# Enzymatic access to the rare ΔUA (1→4) Glc 3, 6, N-sulfated heparin disaccharide, implications for heparin quality control

**DOI:** 10.1101/2024.09.26.615151

**Authors:** T. K. Kandola, C. J. Mycroft-West, M. L. Andrade De Lima, A. Turner, C. Gardini, E. Urso, G.J. Miller, A. Bisio, E. A. Yates, M. Guerrini, L. Wu, M. A. Skidmore

**Affiliations:** Centre for Glycoscience Research and Training, Keele University, Newcastle-under-Lyme, Staffordshire, ST5 5BG, United Kingdom; The Rosalind Franklin Institute, Harwell Science and Innovation Campus, Oxfordshire, OX11 0QX, United Kingdom; Istituto di Ricerche Chimiche e Biochimiche ‘G. Ronzoni’, Via G. Colombo 81, 20133, Milan, Italy; Department of Biochemistry and Systems Biology, Institute of Structural, Molecular and Integrative Biology, University of Liverpool, Crown Street, Liverpool, L69 7ZB, United Kingdom; Division of Structural Biology, Nuffield Department of Medicine, University of Oxford, The Wellcome Centre for Human Genetics, Oxford, OX3 7BN, United Kingdom

## Abstract

The sulfated glycosaminoglycan heparin is the most commonly used pharmaceutical anticoagulant worldwide. Heparin, which is extracted primarily from porcine sources, has a complex heterogeneous structure, resulting in a highly variable pharmaceutical product susceptible to contamination. As a by-product of the food industry, heparin is also limited by production capacity, giving rise to concerns that demand will outstrip supply. The anticoagulant activity of heparin derives principally from the AGA*IA pentasaccharide sequence, containing a rare 3-*O*-sulfated glucosamine, which binds and activates antithrombin. Analytical heparin digestion by the widely used *Pedobacter heparinus* lyases has limited activity in regions of 3-*O*-sulfation, rendering these enzymes poorly suited to study anticoagulant sequences. Here, we provide structural and functional characterization of a *Bacteroides eggerthii* lyase that exhibits highly efficient heparin depolymerization, with specificity distinct to *P. heparinus*. Using a panel of biophysical and structural techniques, we demonstrate that *B. eggerthii* lyase effectively liberates the rare GA* disaccharide, a key indicator of anticoagulant potential, from the defined heparin pentasaccharide fondaparinux. We envision superior cleavage by *B. eggerthii* lyases will enable the future quantitative, direct detection of anticoagulant relevant 3-*O*-sulfated sequences, delivering complementary structural information to existing analytical methods, with clear utility for pharmaceutical quality control workflows.

## Main

Heparin is a highly sulfated glycosaminoglycan (GAG) polysaccharide, widely used as an anticoagulant for the treatment and prophylaxis of thrombotic disorders (Hogwood *et al*., 2023; Hemker, 2016). Heparin, a product of mast cells, exerts anticoagulant properties through the inhibition of blood coagulation proteases (primarily factor IIa (thrombin) and factor Xa), by enhancing the protease inhibition activity of antithrombin (AT) (Lindahl *et al*. 1979; Choay *Et al.*, 1980, Rosenberg and Lam, 1979). Heparin is also structurally related to another member of the GAG family, heparan sulfate (HS) (Menenghetti *et al*., 2015) (**Fig. 1**), which is synthesized by virtually all metazoan cell types, and plays important extracellular functions in cell signaling, differentiation, migration, adhesion, and endocytosis (Ravikumar *et al*., 2020; Collins and Troeberg, 2019; Guimond and Turnbull, 1999; Christianson and Belting, 2014).

**Figure 1.**
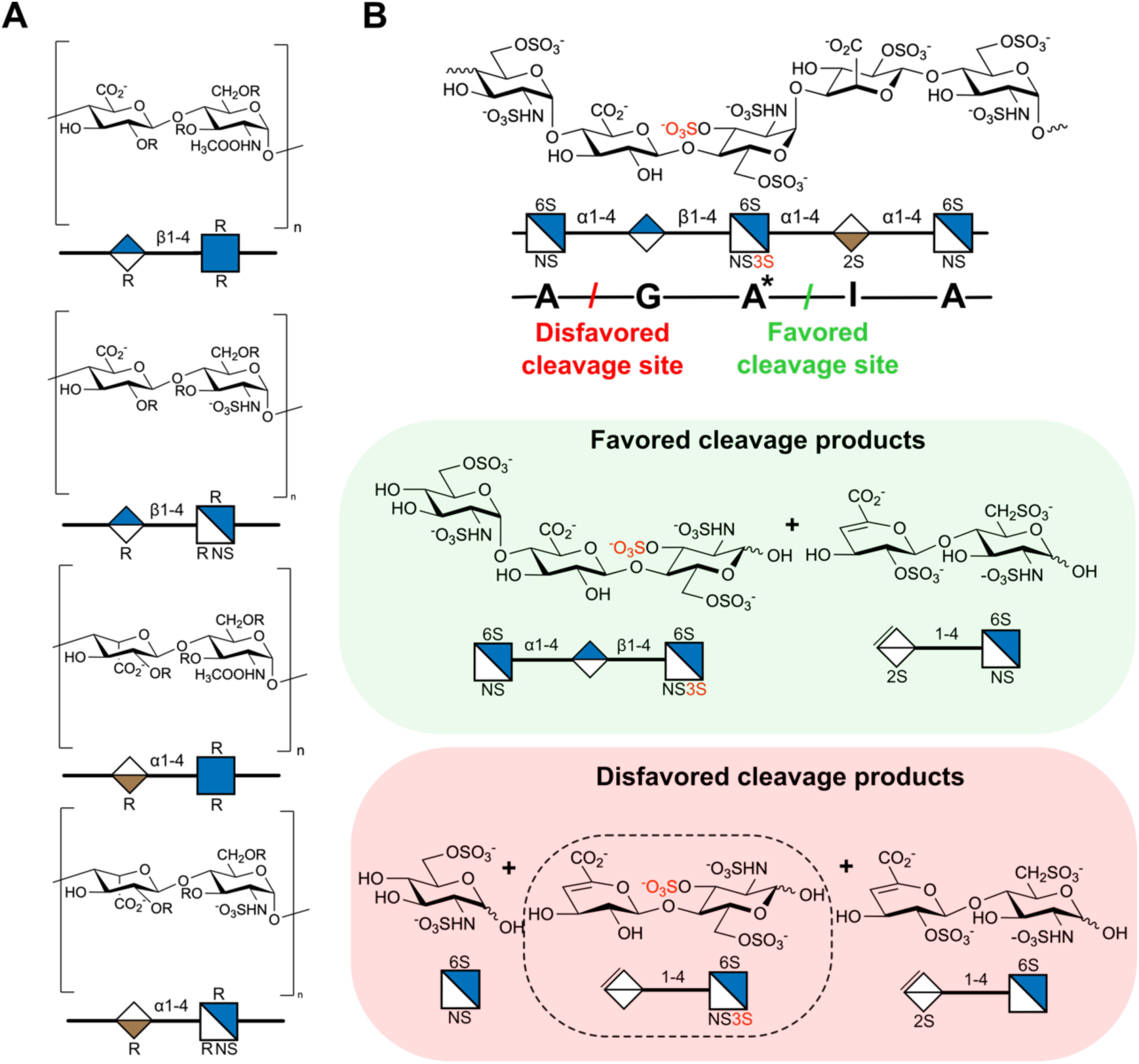
**A**. Major disaccharide repeating units contained within heparin and heparan sulphate (HS) polysaccharide chains, where heparin is composed primarily of the repeating backbone –IdoA–GlcN– and HS – GlcA–GlcN– **B**. Cardinal pentasaccharide motif, attributed to anticoagulant activity of heparin, –GlcNS6S–GlcA– GlcNS3S6S–IdoA2S–GlcNS6S (AGA*IA). **C.** Cleavage of heparin pentasaccharide motif occurs with preference for the AGA*^↓^IA site, while cleavage at A^↓^GA*IA is inefficient.

Structurally, heparin and HS are linear chains of repeating disaccharide units, comprising glucosamines (GlcNAc or GlcNS) 1→4 linked to uronic acids (GlcA or IdoA), further modified by variable sulfation on 2-*O* of IdoA, 6-*O* of GlcNS, and (rarely) 3-*O* of GlcNS (Fig. 1A). Canonically, anticoagulant activity of heparin is mediated by a –GlcNS6S–GlcA–GlcNS3S6S– IdoA2S–GlcNS6S– pentasaccharide motif (AGA*IA, where * denotes 3-*O*-sulfate; Fig. 1B), which binds AT with high affinity (∼ 3 nM), enhancing its protease inhibitory activity (Olson *et al.,* 1992, Björk and Lindahl, 1982, Lima *et al*., 2013). 3-*O*-sulfation enhances AT binding of AGA*IA by 1000-fold compared to AGAIA (Atha *et al*.,1985) and is thus a key modification for anticoagulation action of this sequence (Chopra *et al*., 2021; Atha *et al*., 1985).

The complexity of heparin renders chemical syntheses challenging and costly (Ding *et al*., 2019). Pharmaceutical heparin is therefore primarily obtained from natural sources, mainly of porcine origin, resulting in high batch-to-batch variability. Estimates place only ∼ 1/3 of porcine heparin sequences as possessing anticoagulant activity (Björk and Lindahl, 1982). Further variability arises when considering species of origin, which has slowed the adoption of non-porcine heparin, exacerbating concerns over global supply (Tovar *et al*., 2016). With the use of bovine heparin in South America, and recent developments regarding potential use in the USA (Al-Hakim *et al*., 2021), improved tools are urgently required to ensure the safety and efficacy of this essential pharmaceutical, and support characterization for quality control. One route to achieve this is through advancing methodologies for the quantification of anticoagulant sequences such as AGA*IA.

One of the primary methods for heparin characterization is disaccharide analysis, in which bulk polysaccharide is depolymerized using β-eliminative lyases, and the resulting disaccharides analyzed chromatographically. Currently, the most widely used lyases for heparin/HS digestion are heparinase I – III from *Pedobacter heparinus* (*Ph*HepI – III). However, these enzymes exhibit very limited cleavage of uronic acids towards the non-reducing end of 3-*O*-sulfated glucosamines (e.g. at A^↓^GA*IA), thus cannot reliably discriminate disaccharides containing important 3-*O* modifications (Fig.1B) (Yamada *et al*., 1993, Zhao *et al*., 2011, Shriver *et al*., 2000). This limitation also hinders access to disaccharide standards containing 3-*O*-sulfation, impacting heparin/HS analysis more widely.

Many *Bacteroides* species have evolved to efficiently process complex carbohydrates, as part of their life cycle within intestinal microbiota. Heparin lyase I from *Bacteroides eggerthii* (*Be*HepI) was previously reported to exhibit reduced specificity regard to the uronic acid identity within unfractionated heparin (Bielik *et al*., 2011, Boyce *et al*., 2022). We thus sought to examine whether *Be*HepI may possess augmented heparin depolymerization characteristics compared to *Ph*Heps, using a panel of biophysical and structural techniques to define enzyme activity.

We expressed and purified *Be*HepI from *E coli*, and confirmed its activity against unfractionated heparin, monitoring increase in λ_abs_ = 232 nm over time, corresponding to the unsaturated alkene chromophore generated by lyase digestion (Fig.2A). Further size exclusion chromatography (SEC) of porcine mucosal heparin, exhaustively treated with *Be*HepI or *Ph*HepI, revealed markedly different digestion profiles. The majority of products produced by *Be*HepI were < 3.6 kDa, compared to a substantial amount of poorly digested higher molecular weight material resulting from *Ph*HepI digest (4.2 - 6.0 kDa) (Fig. 2B). The overall broader spread of oligosaccharides produced by *Be*HepI digest vs. *Ph*HepI suggested increased bioactivity by the former towards structural motifs present within this heparin formulation.

**Figure 2.**
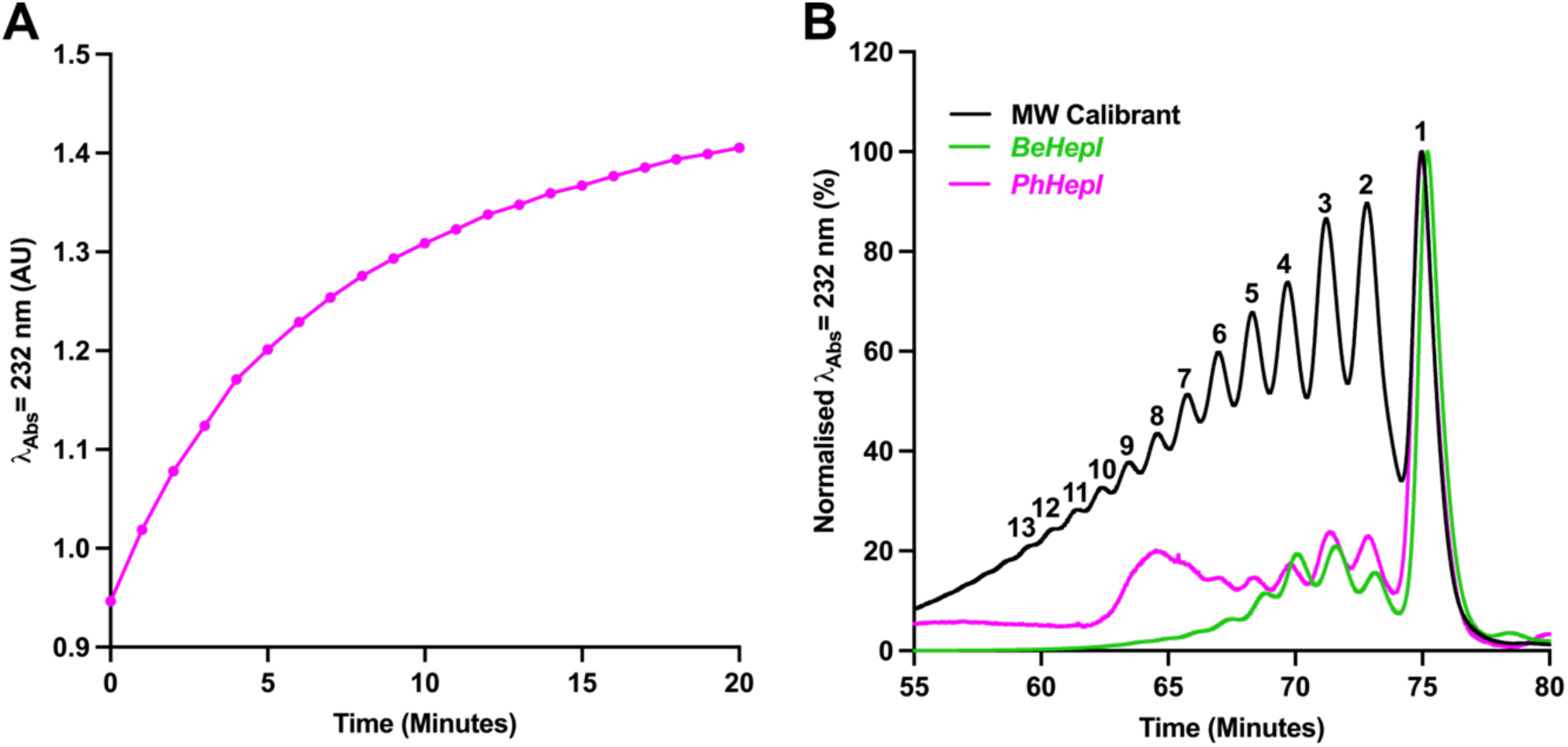
A. Digestion of unfractionated heparin by *Be*HepI digest, monitoring increase in λ_Abs_ = 232 nm **B**. Size exclusion chromatography of unfractionated heparin digested exhaustively with heparinase bacterial lyase enzymes, *Be*HepI (green) or *Ph*HepI (magenta). Molecular weight calibrant (Black; kDa); 1 = 0.6, 2 = 1.2, 3 = 1.8, 4 = 2.4, 5 = 3.0, 6 = 3.6, 7 = 4.2, 8 = 4.8, 9 = 5.4, 10 = 6.0, 11 = 6.6, 12 = 7.2, 13 = 8.4.

The heterogeneity of unfractionated heparin hinders analysis of enzyme specificities. To precisely characterize *Be*HepI cleavage, we thus turned to the defined pentasaccharide fondaparinux sodium (Arixtra®), a fully synthetic heparin mimetic corresponding to reducing end *O*-methylated AGA*IA. Digestion of fondaparinux by *Ph*HepI yielded disaccharide and trisaccharide fragments corresponding to AGA*^↓^IA, consistent with previous reports (Yamada *et al*., 1993, Zhao *et al*., 2011, Shriver *et al*., 2000). Notably, a large abundance of ion m/z 456.97 (z = −2), consistent with the trisaccharide product AGA* (m/z assignment; A3,5,0), indicated that cleavage of fondaparinux sodium by *Ph*HepI is notably inefficient at A^↓^GA*IA sites (**Table S1**). In stark contrast, *Be*HepI produced a distinct digestion pattern, with products corresponding to digestion at both A^↓^GA*^↓^IA positions. Firstly, ion of m/z 337.99 (−1), assigned as A1,2,0, indicates the liberation of the glucosamine monosaccharide (A) from the non-reducing end (NRE) of fondaparinux sodium. Ions of m/z 575.96 (−1) (assignment; ΔU2,3,0) and m/z 589.98(−1) (assignment; ΔU2,3,0-OMe) were also detected, which are consistent with the release of both GA* and IA-OMe disaccharides, respectively (**Fig. 3, Table S2**). Furthermore, the ion with m/z 705.10 (−1), could be assigned as ΔU2,3,0 GA*+DBA, and therefore likely corresponds to liberation of disaccharide GA*+DBA. The marked increase in abundance of the m/z 575.96 and m/z 705.10 (−1) disaccharide ions in comparison to *Ph*HepI, is therefore consonant with the liberation of the GA* disaccharide by *Be*HepI. This can clearly be seen in the EIC corresponding to BPC peak 4, between *Be*HepI and *Ph*HepI digests *Be*HepI (**Fig. 3, Table S1, Table S2**). Moreover, elevated abundance of the m/z 337.99 (−1) monosaccharide ion and the corresponding reduced levels of the m/z 456.9741 (−2) trisaccharide ion between the *Be*HepI and *Ph*HepI digests, further supports enhanced cleavage of A^↓^GA*IA by *Be*HepI (**Fig. 3, Table S1, Table S2**).

**Figure 3.**
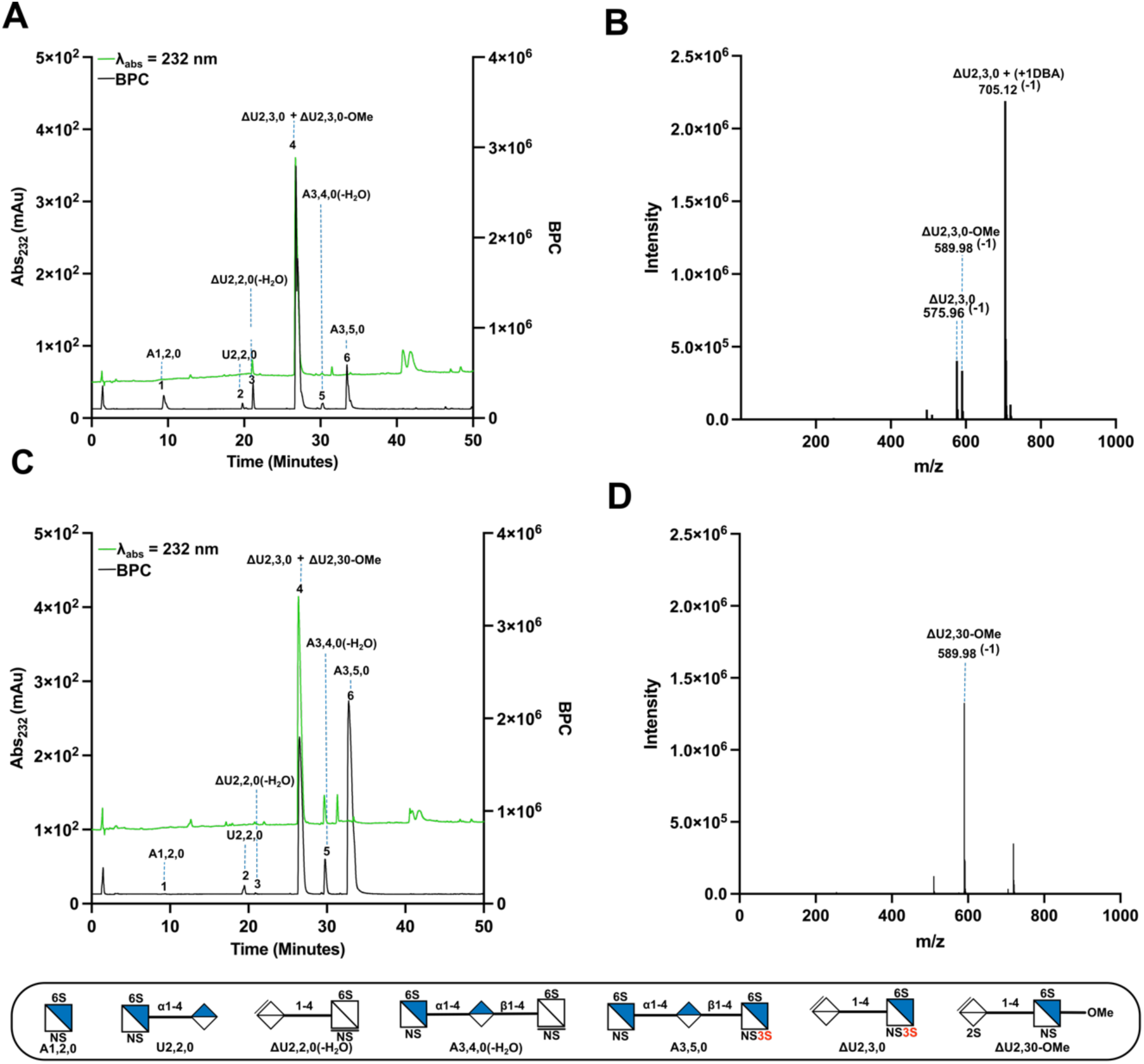
A. LC-MS base peak chromatogram (BPC) (black) and λ_Abs_ = 232 nm (green) of major fondaparinux digestion fragments produced by *Be*HepI. **B.** Mass spectrum corresponding to BPC, peak4 from *Be*HepI cleavage; m/z 575.96 (−1) (ΔU2,3,0; GA*), m/z 589.98(−1) (ΔU2,3,0-OMe, IA-OMe) and m/z 705.1(−1) (ΔU2,3,0 (+1DBA), GA* +1DBA). **C**. LC-MS BPC and λ_Abs_ = 232 nm of major fondaparinux digestion fragments produced by *Ph*HepI. **D**. Mass spectrum corresponding to peak 4 from *Ph*HepI cleavage, m/z 589.98(−1) (ΔU2,3,0-OMe; IA-OMe). Assignment nomenclature from the nonreducing end (NRE) corresponds to ΔU or U, indicating the 4,5-unsaturated uronic acid and saturated uronic acid residues, respectively or A for glucosamine units. Subsequent numbering, in order, represents the number of monosaccharide residues, sulfate groups and acetyl constituents.

To further confirm the presence of digested GA* and IA disaccharides from *Be*HepI digestion of fondaparinux, 1-D ^1^H and 2-D ^1^H-^13^C NMR spectroscopy was conducted. We observed the appearance of alkene proton signals at 5.86 and 5.89 ppm following exhaustive *Be*HepI treatment for 48 hours, indicating two distinct unsaturated products from cleavage (**Figs. 4 & S1**). Partial digestion experiments on replicate samples showed the 5.86 ppm peak to appear first, followed by the 5.89 ppm peak, suggesting differences in cleavage efficiencies for their corresponding glycosidic bonds (**Fig. S1**). Cross-referencing to the anomeric peaks for IdoA and GlcA in undigested fondaparinux sodium (5.07 ppm and 4.52 ppm respectively) showed rapid loss of the IdoA peak, followed by slower loss of the GlcA peak, which required extended digestion conditions to process (**Fig. S1**). Taken together, our data suggest that *Be*HepI efficiently digests glycosidic bonds to both GlcA and IdoA in the context of fondaparinux sodium, albeit with increased efficiency for the bond to IdoA.

**Figure 4.**
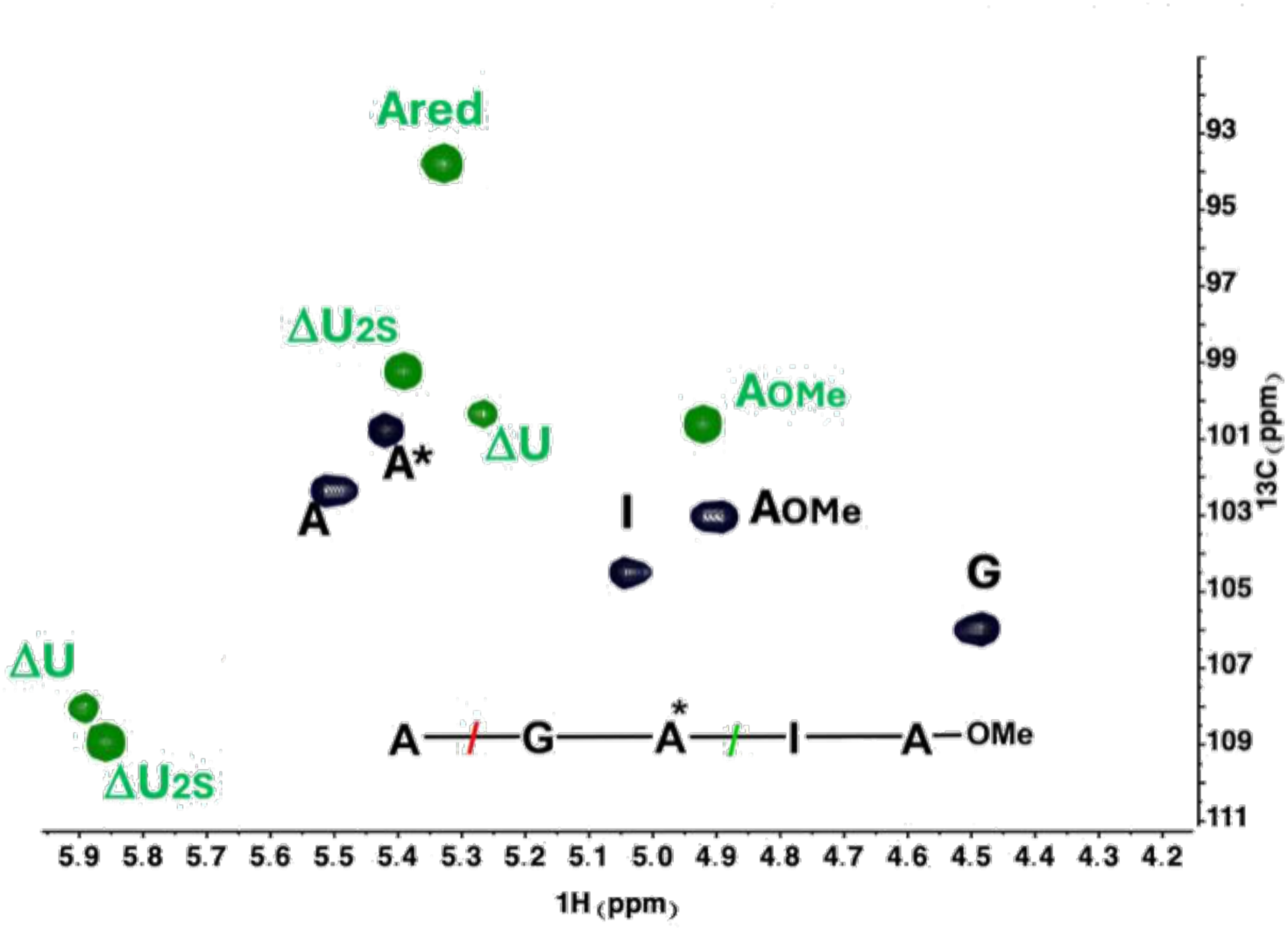
^1^H-^13^C HSQC NMR spectrum of fondaparinux sodium after digestion with *Be*HepI.

Finally, to directly confirm the digestion preference of *Be*HepI, we determined X-ray crystal structures of the enzyme in both apo form (PDB 9GRW) and, after application of fondaparinux (PDB 9GV9). The determined crystal structures were isomorphous and processed to 1.7 Å and 1.9 Å for the apo and liganded structures respectively (**Fig. S2, Table S3**). Two molecules of *Be*HepI were present in each asymmetric unit of the structures, with a relatively small contact interface (∼ 819 Å^2^) consistent with a crystallographic packing interaction.

The 3-dimensional structure of unliganded *Be*HepI resembled that of the related and previously reported HepI from *Bacteroides thetaiotamicron* (*Bt*HepI; PDB 3IKW; RMSD 1.9 Å over 28 −394 amino acids; 84% identity) (Han *et al*., 2009), and comprises a central β-jelly roll fold (residues 28 - 111, 139 - 175, 242 - 391) with an extended Ω-loop (residues 112 - 138) and ‘thumb’ domain (residues 176 - 241). The inner antiparallel β-sheet of the β-jelly roll and the thumb domain contribute to the formation of an extended, positively charged substrate binding cleft containing a central groove and overhanging lid (Fig. 5). The thumb and the β-jelly roll domains are connected through the involvement of the sidechain oxygen atoms of Glu240 and Asp364, the main chain carbonyl oxygens of Trp266 and Asn363, and two water molecules, *via* an octahedral coordinated Ca^2+^ ion (**Figs. 5A & S3**).

**Figure 5:**
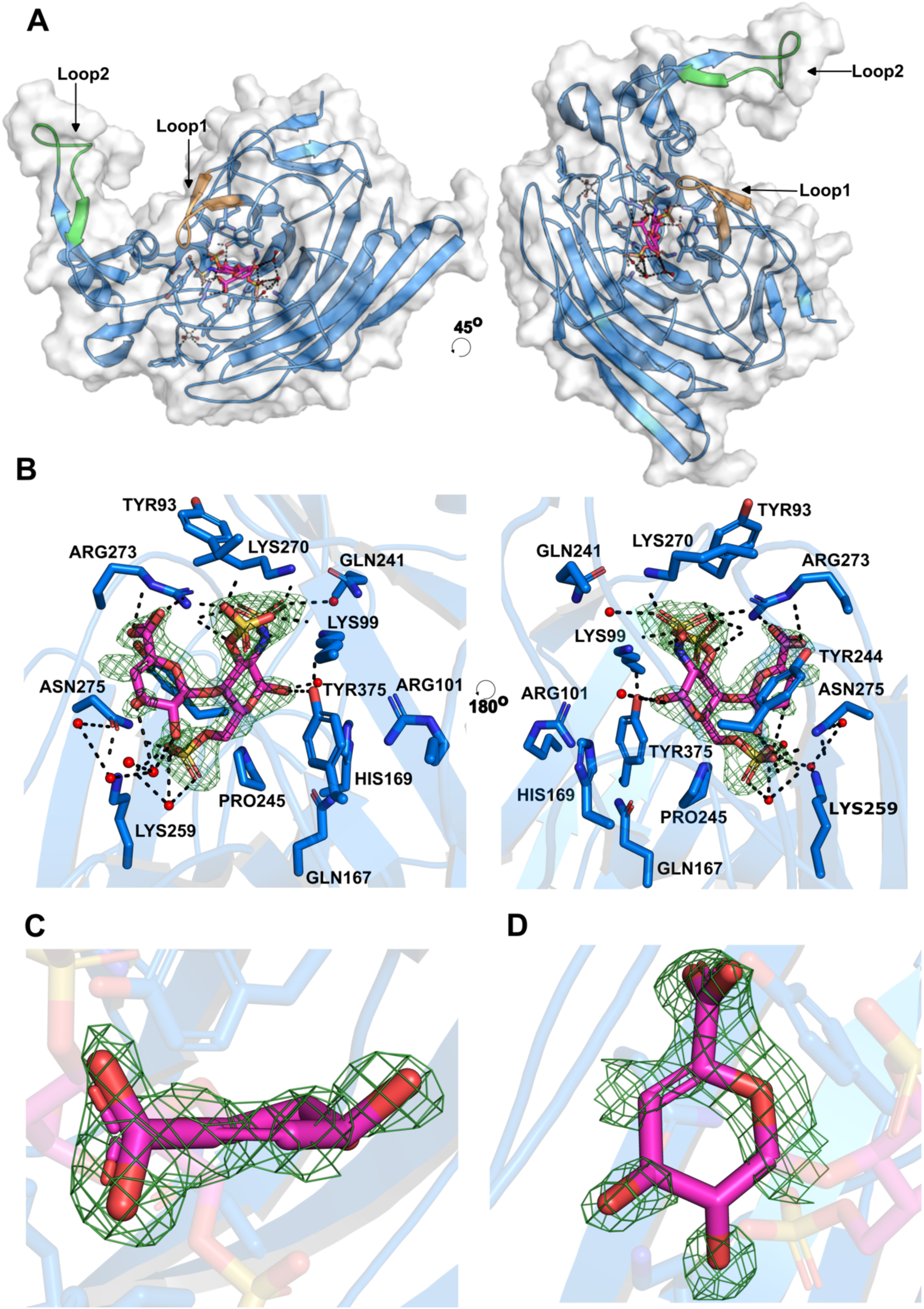
**A** Structure of *Be*HepI bound to GA* disaccharide product. **B**. Active site views in complex with GA* disaccharide product in the −2 and −2 subsites. **C & D**. Close-up view of uronic acid unsaturated C4-C5 bond resulting from β-elimination cleavage, assuming a half-chair conformation. Electron density for ligand is REFMAC σ_A_-weighted mFo-DFc, contoured to 2 σ.

The structure of *Be*HepI after the addition of fondaparinux sodium was broadly similar to the *apo* structure but contained clear additional density within the enzyme active site cleft, which was well modelled by the GA* unsaturated disaccharide (real-space correlation coefficient 0.93), the expected 3*-O*-sulfated product after cleavage (Fig. 5). The GA* disaccharide occupied the –1, –2 subsites of the active site cleft (David, Wilson and Henrissat, 1997), with the reducing-end hydroxyl hydrogen-bonded to Tyr375. Further hydrogen-bond interactions were observed between the *N*-sulfate of GlcNS3S6S, the side chain nitrogen atom of Lys270 and the backbone nitrogen atoms of Gly223 and Arg253. The 3-*O*-sulphate of GlcNS,3S,6S formed hydrogen-bonds with the backbone nitrogen atom of Tyr93 and the side chain nitrogen atom of Lys9. While the 6-*O*-sulphate of GlcNS3S6S formed hydrogen-bonds with the side chain nitrogen atom of Asn75 and Lys259. The carboxyl group of Δ4,5UA was also observed to form hydrogen-bonds with the sidechain of Arg273 and Tyr244.

Whilst binding of GA* to the *Be*HepI –1 and –2 subsites implies enzymatic attack at AGA*^↓^IA, rather than A^↓^GA*IA, we also observed clear evidence supporting cleavage of the bond to glucuronic acid. Notably, electron density around the non-reducing end of the bound disaccharide adopted a clear planar geometry around C4–C5, consistent with the presence of an unsaturated double bond following lyase cleavage (Fig. 5). Electron density around the C4–C5–(C6)–O5 bonds was also stronger compared to other areas of the ring, indicative of an extended π-system encompassing the alkene, endocyclic oxygen, and C6 carboxylate (**Figs. 5 & S4**). Conversely, some minor density around O4 of the uronic acid was visible, likely corresponding to a small amount of uncleaved AGA* trisaccharide (Fig. 5). Taken together, these structures are consistent with LC-MS and NMR data and suggest initial rapid attack of the bond to iduronic acid, followed by slower cleavage of the bond to glucuronic acid, ultimately resulting in dual A^↓^GA*^↓^IA digest. The improved efficiency of A^↓^GA*IA cleavage may reflect underlying affinities of *Be*HepI for different sulfation or epimerization patterns. However, A^↓^GA*IA cleavage would also demand only 1 sugar, the terminal GlcNS6S, within the ‘minus’ region of the *Be*HepI substrate cleft, which may be detrimental for affinity. As with other lyases, further digest studies using longer defined oligosaccharides will be required to fully deconvolute specificity patterns for *Be*HepI.

No experimental structure currently exists for *Ph*HepI, limiting our ability to directly examine its limited tolerance for anticoagulant heparin sequences. The AlphaFold prediction model of *Ph*HepI exhibits a similar 3-dimensional structure to that of *Be*HepI (and *Bt*HepI), with the notable absence of two loops, around the substrate binding site; Thr89 - Glu100 (loop 1) and Leu213 - Gly232 (loop 2). Importantly, residues Tyr93 and Lys99 of *Be*HepI, present in loop 1, make H-bonding and electrostatic interactions with the 3-*O*-sulfate of the disaccharide product, which are likely to improve *Be*HepI affinity for this structural feature. The presence of loops 1 and 2 also contributes to a more positive and extended electrostatic patch within the active site of *Be*HepI compared to the *Ph*HepI model, consistent with improved activity of *Be*HepI against more negatively charged oligosaccharide substrates (**Figs. 5 & S5 - S6**)

In summary, we have defined the activity *Be*HepI, which displays distinct substrate specificity compared to the widely used *Ph*Heps. Of particular interest is the efficient cleavage of *Be*HepI at uronic acids immediately adjacent to 3*-O*-sulfated glucosamines, a position that is poorly recognized by *Ph*HepI. Such sequence specificity renders *Be*HepI well-suited to the analytical digestion of heparin, as well as for the efficient preparation of standards containing 3-*O*-sulfation from bulk polysaccharide. Our work has clear implications for the characterization of anticoagulant heparin motifs and will facilitate improved quality control and standardization of this vital global pharmaceutical.

## Methods and Materials

### Expression and purification of heparinase I from Bacteroides eggerthii

The heparinase I gene from *Bacteroides eggerthii* (WP_004294077.1), lacking the N-terminal leader sequence (21 - 394), was inserted into the pET-22b(+) plasmid (Merck, UK) and transformed into OverExpress™ C41 (DE3) pLysS (Imaxio SA, France) chemically competent cells as per the manufacturer’s instructions. Transformed cells were grown under agitation at 37°C in 10 mL Miller’s LB Broth (LB; Melford, UK) containing 100 µg.mL^-1^ and 50 µg.mL^-1^ chloramphenicol. After 16 hours, 1L of Terrific Broth (TB; Melford, UK) supplemented with 100 µg.mL^-1^ and 50 µg.mL^-1^ chloramphenicol (Apollo Scientific, UK) was inoculated with the pre-culture and grown under agitation at 37°C until the O.D._600_ reached 0.6 - 0.8. The culture was then cooled to 16°C and expression induced with 1 mM IPTG (Apollo Scientific, UK), before being incubated under agitation for a further 16 hours. The culture was harvested by centrifugation at 6,000g for 15 minutes, the supernatant discarded, and the cell pellet resuspended in 25 mM Tris-HCl, 500 mM NaCl, 5 mM CaCl_2_, 20 mM imidazole pH 7.4, containing cOmplete^TM^ protease inhibitor cocktail (Merck, UK) and 120 Kunits DNase (Merck, UK), before being subjected to two rounds of cell disruption at 4°C, 30 kpsi (CF1-AE; Constant Systems, UK). The culture was subsequently centrifuged at 16,000g for 30 minutes, 4°C and the supernatant collected, before being loaded onto a 5 mL HisTrap FF column (Cytiva, UK) at 5 mL.min^-1^, which had been equilibrated in 25 mM Tris-HCl, 500 mM NaCl, 5 mM CaCl_2_, 20 mM imidazole, pH 7.4. The loaded column was washed with 5 column volumes of 25 mM Tris-HCl, 500 mM NaCl, 5 mM CaCl_2_, 20 mM imidazole, pH 7.4, then bound material eluted with a linear gradient of 0 – 100% 25 mM Tris-HCl, 500 mM NaCl_2_, 5 mM CaCl_2_, 200 mM imidazole over 20 column volumes. Fractions containing protein with a molecular weight corresponding to BeHepI were determined by SDS-PAGE, pooled and concentrated using a 10 kDa molecular weight cut-off centrifugal filter (Vivaspin® Turbo 15; Sartorius, UK) prior to being applied to a S75 column pre-equilibrated in 25 mM Tris, 100 mM NaCl, 5 mM CaCl_2_ pH 7.4. Elution was performed in the same buffer, isocratic at 1 mL.min^-1^. BeHepI containing fractions, as determined by SDS-PAGE, were pooled, diluted 10-fold into 20 mM MES, 5 mM CaCl_2_, pH 6.0 and then applied to a Resource S cation exchange column (Cytiva; UK) at 2 mL.min^-1^. The column was washed over 10 column volumes with 20 mM MES, 5 mM CaCl_2_ before bound material was eluted using a linear gradient of 0 - 100% 20 mM MES, 5 mM CaCl_2_, 1.5 M NaCl, pH 6.0. Fractions containing heparinase I at > 95% purity, as determined by SDS-PAGE, were subsequently concentrated and buffer exchanged into 20 mM Tris, 100 mM NaCl, 5 mM CaCl2 pH 8.0 (3 times the original volume) using a 10 kDa molecular weight cut-off centrifugal filter (Vivaspin® Turbo 15; Sartorius, UK), before flash-freezing in liquid nitrogen and storage at −80°C until required.

### Saccharide digestion by heparinase I

Fondaparinux (Sigma-Aldrich, SML1240) and unfractionated sodium heparin (Celsus, OH, USA) were subjected to exhaustive depolymerisation by *BeHepI* or *PhHepI* in a buffer containing 100 mM sodium acetate and 0.1 mM calcium acetate, pH 7.0 for 48 hours at 37°C under constant agitation. An equal quantity of enzyme was added at time points, 0, 1, 2 and 24 hours, totalling 160 mIU. After 48 hours the reaction mixture was heat inactivated (95°C for 2 minutes), filtered (0.22 μm cellulose acetate; LLG labware, UK) and analysed by LC-MS and NMR.

### Gel permeation chromatography

Oligosaccharide profiling was performed using gel permeation chromatography (TSK G3000SW, 7.8mm ID x 30 cm, 5.0 μm and TSKG2000SW, 7.8mm ID x 30 cm, 5.0 μm; columns connected in series; Tosoh Bioscience GmbH, Germany) with a mobile phase of 0.3 M sodium sulphate, on a high-pressure liquid chromatography system equipped with in line UV detection at λ_abs_ = 232 nm (Agilent, UK). Isocratic elution was conducted at 300 μL.min^-1^ and the system was previously calibrated with the Heparin Low-Molecular-Mass for Calibration CRS (EDQM, France).

### Mass spectrometry

Mass spectrometric analyses were performed on lyase digested samples using a reverse phase HPLC system (Platin Blue, Knauer) coupled inline to a Bruker ESI-Q-TOF Impact II Daltonics system (Bruker, Italy). The chromatographic separation was achieved by C-18 Kinetex column (100 x 2.1 mm, 2.6 µm, 100Å, Phenomonex, Italy), applying a multi-step gradient elution method with Phase A (10 mM Dibutylamine (≥ 99.5%), 10 mM ethanol; Merck, Italy) and Phase B (10 mM DBA, 10 mM ethanol, in LC-MS grade methanol; Merck, Italy) with a flow rate of 0.15 ml.min^-1^ at 35°C. 8 µg of digested sample was injected for each heparinase I digest onto the LC-MS setup, with MS parameters for analysis of the digested products as follows; ESI in negative ion mode; capillary voltage + 3,500V; nebulizer pressure 1.8 bar; dry gas flow 7.0 l.min^-1;^ dry gas temperature 200°C, and a m/z scan range of 140 – 2,500.

### Nuclear Magnetic Resonance

1-D (^1^H) and 2-D (^1^H-^13^C) Heteronuclear Single Quantum Coherence (HSQC), 2-D COSY and 2-D TOCSY spectra (**Figs. S7 - 9**) of the exhaustive fondaparinux sodium digests were recorded using a Bruker Avance NEO 600 MHz (Bruker, UK) spectrometer fitted with a BBO ^1^H-^19^F cryoprobe (Bruker, UK) at 298 K. Nuclear Magnetic Resonance spectra of partial fondaparinux sodium digests were recorded on a Bruker Avance 400 spectrometer equipped with a BBFO probe (Bruker, UK). The chemical shift data for each signal are given as δ in units of parts per million (ppm) relative to tetramethylsilane, where δ = 0.00 ppm. Both 1-D and 2-D NMR spectra were collected using standard Bruker pulse sequences.

### Crystallisation of heparinase I from *Bacteroides eggerthii*

BeHepI in 20 mM Tris, 100 mM NaCl, 5 mM CaCl_2_, pH 8.0 was concentrated to 10 mg.mL^-1^ using 10 kDa molecular weight cut-off spin filter (Vivaspin® Turbo 15; Sartorius, UK) and screened using commercially available screens, PACT premier (Molecular Dimensions, UK) and JCSG Plus (Molecular Dimensions, UK), by sitting drop vapour diffusion at 20°C, with an equal protein:well ratio of 150 nL. Well-diffracting crystals were obtained in a precipitant of 2M (NH_4_)_2_SO_4_, 0.1 M CH_3_CO_2_Na, pH 5.0.

Ligand complexes with fondaparinux sodium (Merck, UK) were obtained by addition of the powdered ligand directly into drops containing crystals, followed by incubation for 5 minutes before harvesting and cryo-cooling crystals. Diffraction data was collected using the i04 beamline of Diamond Light Source (Harwell, UK), equipped with an Eiger2 XE 16M pixel detector (Dectris AG, Switzerland), at a wavelength of 0.9763 Å. Liganded and apo datasets were processed with the autoPROC+ STARANISO pipeline and xia2 dials, respectively (Global Phasing, UK). Phasing was subsequently solved by molecular replacement with MOLREP (Vagin and Teplyakov, 2010), using heparin lyase I from *B. thetaiotaomicron* (PDB 3IKW) as a search model. The determined structure was improved by iterative rounds of manual model building with COOT (Emsley and Cowtan, 2004), alongside maximum-likelihood refinement with REFMAC5 (Murshudov *et al*., 2011). Ligand coordinates and dictionaries were defined using JLigand (Lebedev *et al*., 2012). Structure figures were generated using Pymol (Schrödinger Inc., NY, USA).

## Abbreviations

BMH: bovine mucosal heparin
BSE: 
DoS: degree of sulfation
GAG: glycosaminoglycan
GalN: D-galactosamine
GlcN: D-glucosamine
HS: heparan sulfate
LMWH: low molecular weight heparin
OSCS: over-sulfated chondroitin sulfate
OMH: ovine mucosal heparin
NMR: nuclear magnetic resonance
PMH: porcine mucosal heparin

## Supplementary tables and figures

**Table S1.** Mass spectrum assignment of the main peaks from the Base Peak Chromatogram of fondaparinux sodium digested with *Ph*HepI.

| Peak | m/z exp | z | M | Assignment | M/z th. | Error (ppm) | Proposed Structure | Linkages involved in the enzymatic mechanism and fragments released (red) |
| --- | --- | --- | --- | --- | --- | --- | --- | --- |
| 1 | 337.9848 | -1 | 339 | A1,2,0 | 337.9857 | 2.6 | A<sub>NS6S</sub> | A→GA*IA |
| 2 | 514.0155 | -1 | 515 | U2,2,0 | 514.0178 | 4.4 | A<sub>NS6S</sub>-G | Second step peeling from A<sub>NS6S</sub>-G-A<sub>NS6S</sub>(-H<sub>2</sub>O) |
| 3 | 477.9947 | -1 | 479 | ΔU2,2,0(-H<sub>2</sub>O) | 477.9967 | 4.1 | ΔU-A<sub>NS6S</sub>(-H<sub>2</sub>O) | First step peeling from ΔU-A<sub>NS3S6S</sub> |
| 4 | 589.9768 | -1 | 591 | ΔU2,3,0-OMe | 589.9797 | 4.9 | ΔU2S-A<sub>NS6S</sub>(OMe) | AGA*→ IA |
| 5 | 407.9904 | -2 | 818 | A3,4,0(-H<sub>2</sub>O) | 407.9912 | 1.9 | A<sub>NS6S</sub>-G-A<sub>NS6S</sub>(-H<sub>2</sub>O) | First step peeling from A<sub>NS6S</sub>-G-A<sub>NS3S6S</sub> |
| 6 | 456.9741 | -2 | 916 | A3,5,0 | 456.9749 | 1.8 | A<sub>NS6S</sub>-G-A<sub>NS3S6S</sub> | AGA*→ IA |

**Table S2.** Mass spectrum assignment of the main peaks from the Base Peak Chromatogram of fondaparinux sodium digested with *Be*HepI.

| Peak | m/z exp | z | M | Assignment | M/z th. | Error (ppm) | Proposed Structure | Linkages involved in the enzymatic mechanism and fragments released (red) |
| --- | --- | --- | --- | --- | --- | --- | --- | --- |
| 1 | 337.9859 | -1 | 339 | A1,2,0 | 337.9857 | 0.6 | A<sub>NS6S</sub> | A→GA*IA |
| 2 | 514.0232 | -1 | 515 | U2,2,0 | 514.0178 | 10.5 | A<sub>NS6S</sub>-G | Second step peeling from A<sub>NS6S</sub>-G-A<sub>NS6S</sub>(-H<sub>2</sub>O) |
| 3 | 477.9971 | -1 | 479 | ΔU2,2,0(-H<sub>2</sub>O) | 477.9967 | 0.8 | ΔU-A<sub>NS6S</sub>(-H<sub>2</sub>O) | First step peeling from ΔU-A<sub>NS3S6S</sub> |
| 4 | 705.1161+<br>589.9803 | -1<br>-1 | 706<br>591 | ΔU2,3,0 (+1DBA)<br>ΔU2,3,0-OMe | 705.1158<br>589.9797 | 0.4<br>1.0 | ΔU-A<sub>NS3S6S</sub><br>ΔU2S-A<sub>NS6S</sub>(OMe) | A→GA*→ IA<br>AGA*→ IA |
| 5 | 407.9924 | -2 | 818 | A3,4,0(-H<sub>2</sub>O) | 407.9912 | 2.9 | A<sub>NS6S</sub>-G-A<sub>NS6S</sub>(-H<sub>2</sub>O) | First step peeling from A<sub>NS6S</sub>-G-A<sub>NS3S6S</sub> |
| 6 | 456.9759 | -2 | 916 | A3,5,0 | 456.9749 | 2.2 | A<sub>NS6S</sub>-G-A<sub>NS3S6S</sub> | AGA*→ IA |

**Supplementary Figure 1.**
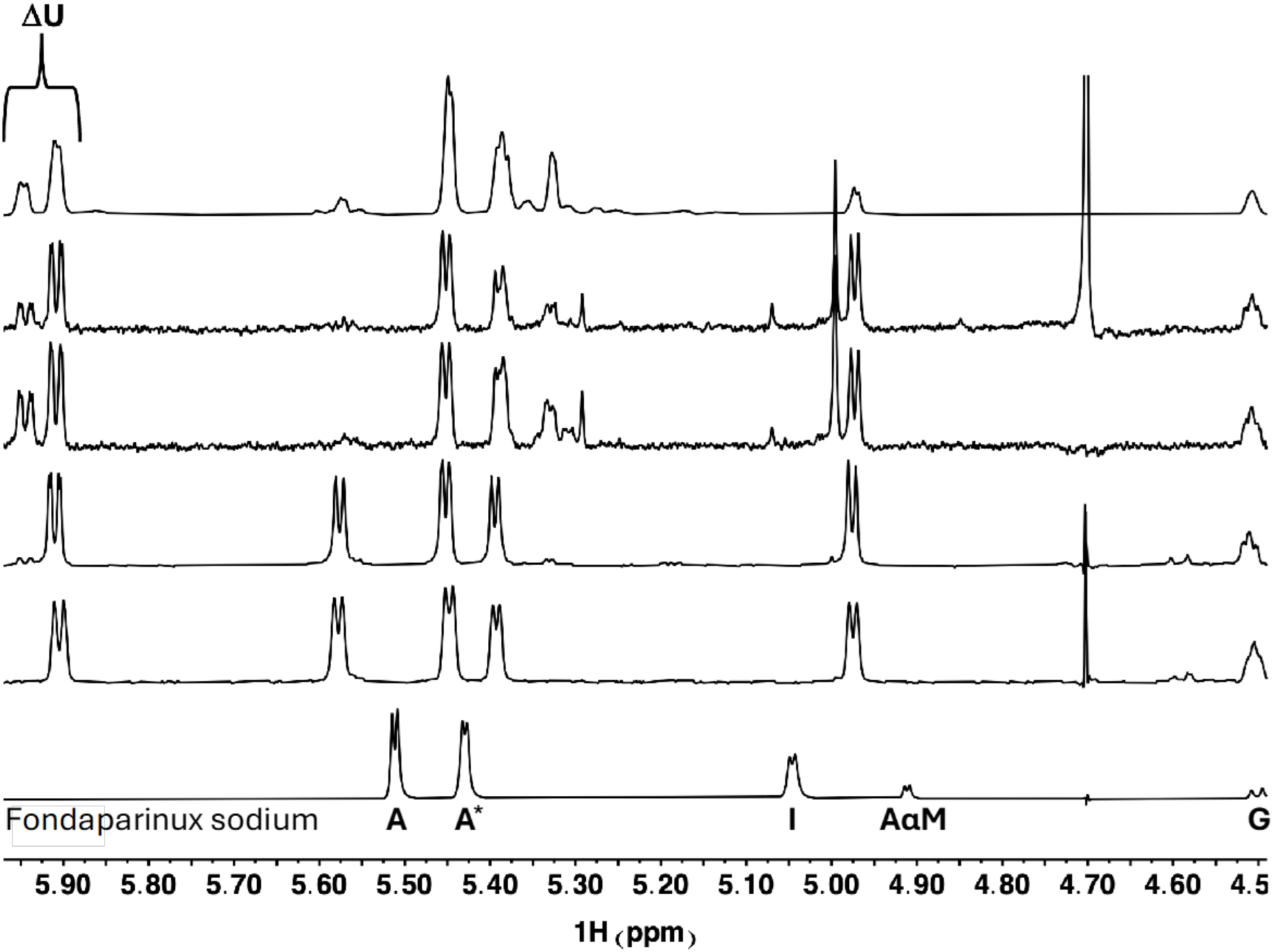
1-D ^1^H NMR spectra of distinct levels of fondaparinux sodium digestion by *Be*HepI, progressing from the undigested substrate (bottom) to the fully digested, end-product (top)

**Supplementary Table 3.**
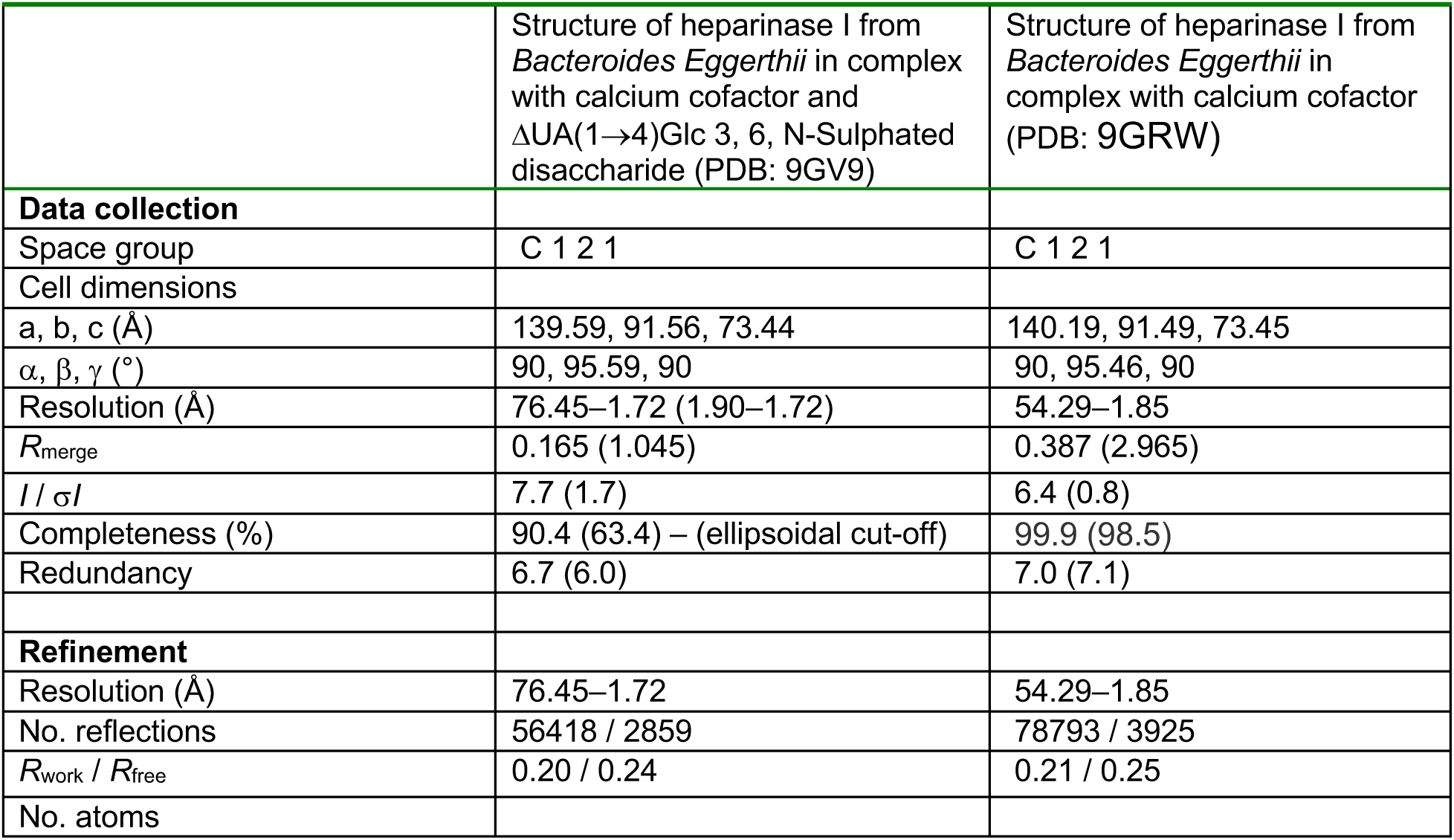
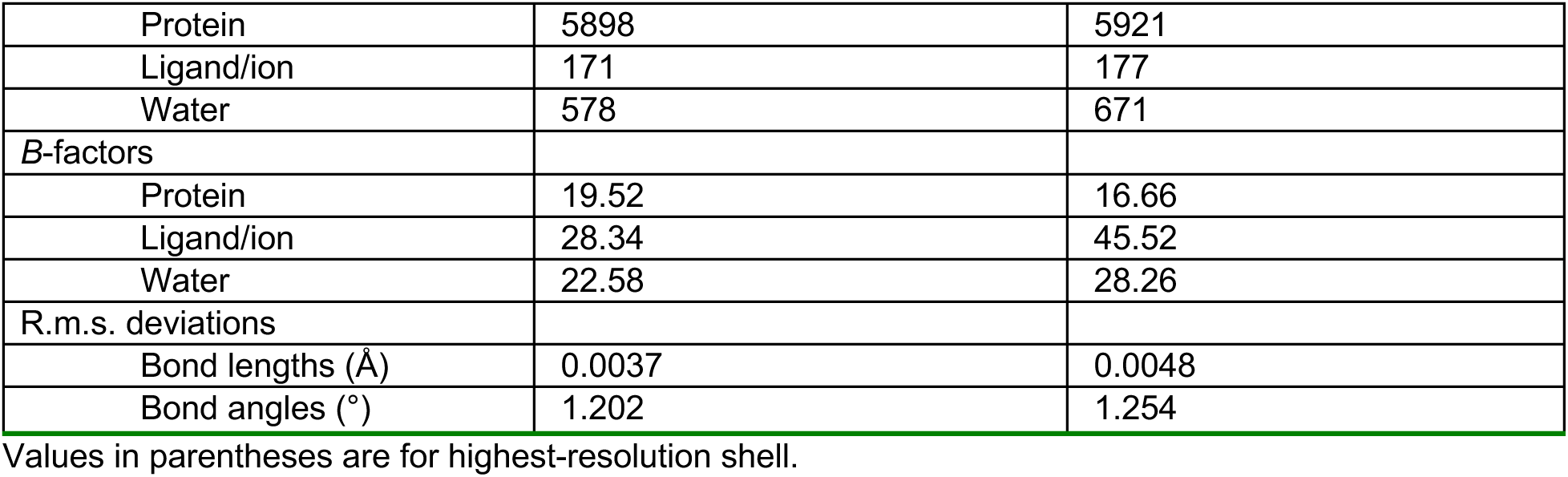
Data collection and refinement statistics (molecular replacement)

**Supplementary Figure 2.**
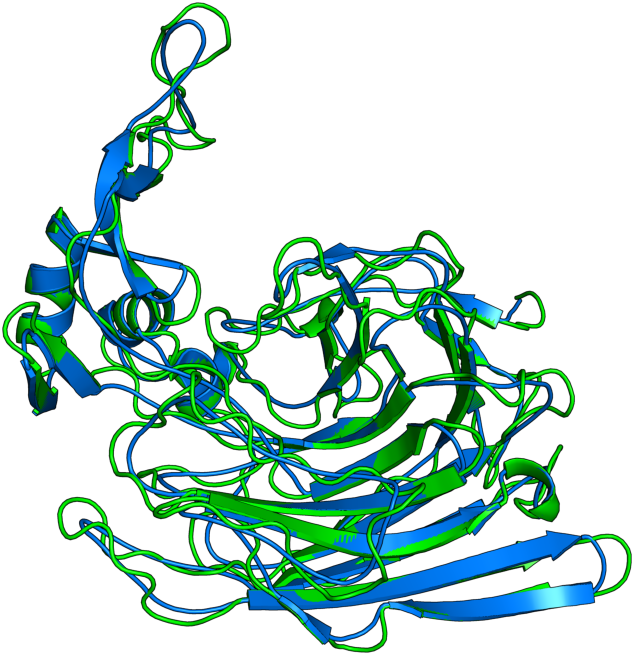
Overlay between apo (green) and ligand structure (blue)

**Supplementary Figure 3.**
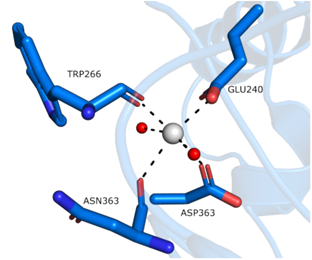
Calcium binding site.

**Supplementary Figure 4.**
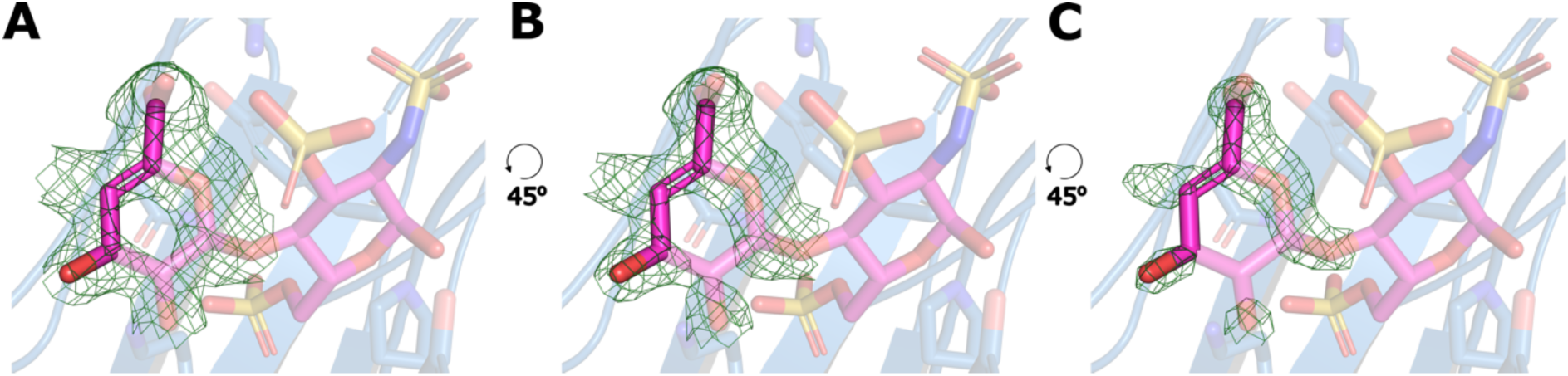
Difference map levels at different weightings **A)** contoured to 1.5 σ **B)** contoured to 2 **C)** contoured to 3.

**Supplementary Figure 5.**
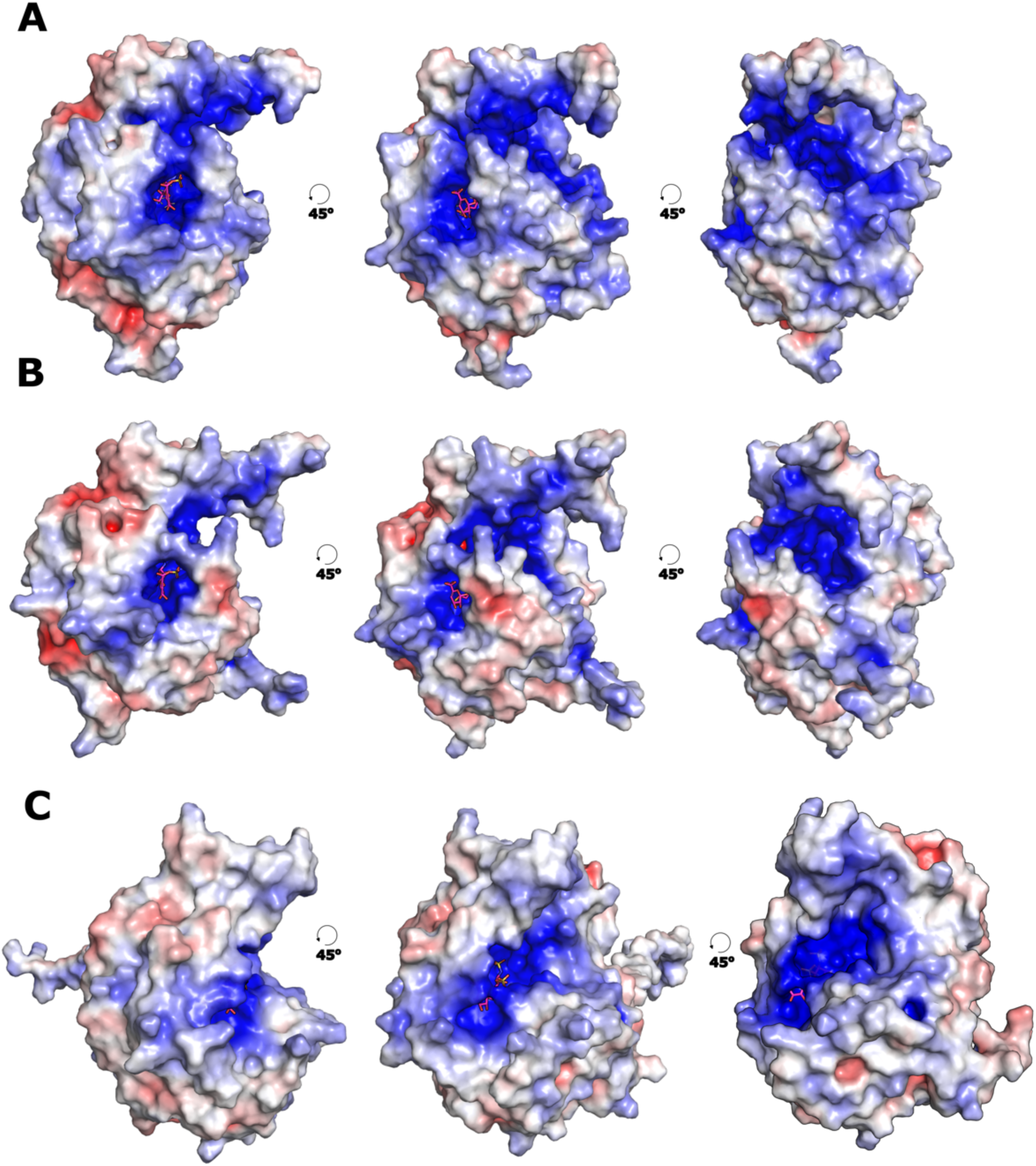
Comparison of the Coulomb electrostatic potential maps of **A.** *Be*HepI, **B**. *Bt*Hep1 and **C.** *Ph*HepI, with an overlay of the fondaparinux digestion product (ΔGlcA (α1→4) Glc 3, 6, NS) binding site from PDB 9GV9.

**Supplementary Figure 6.**
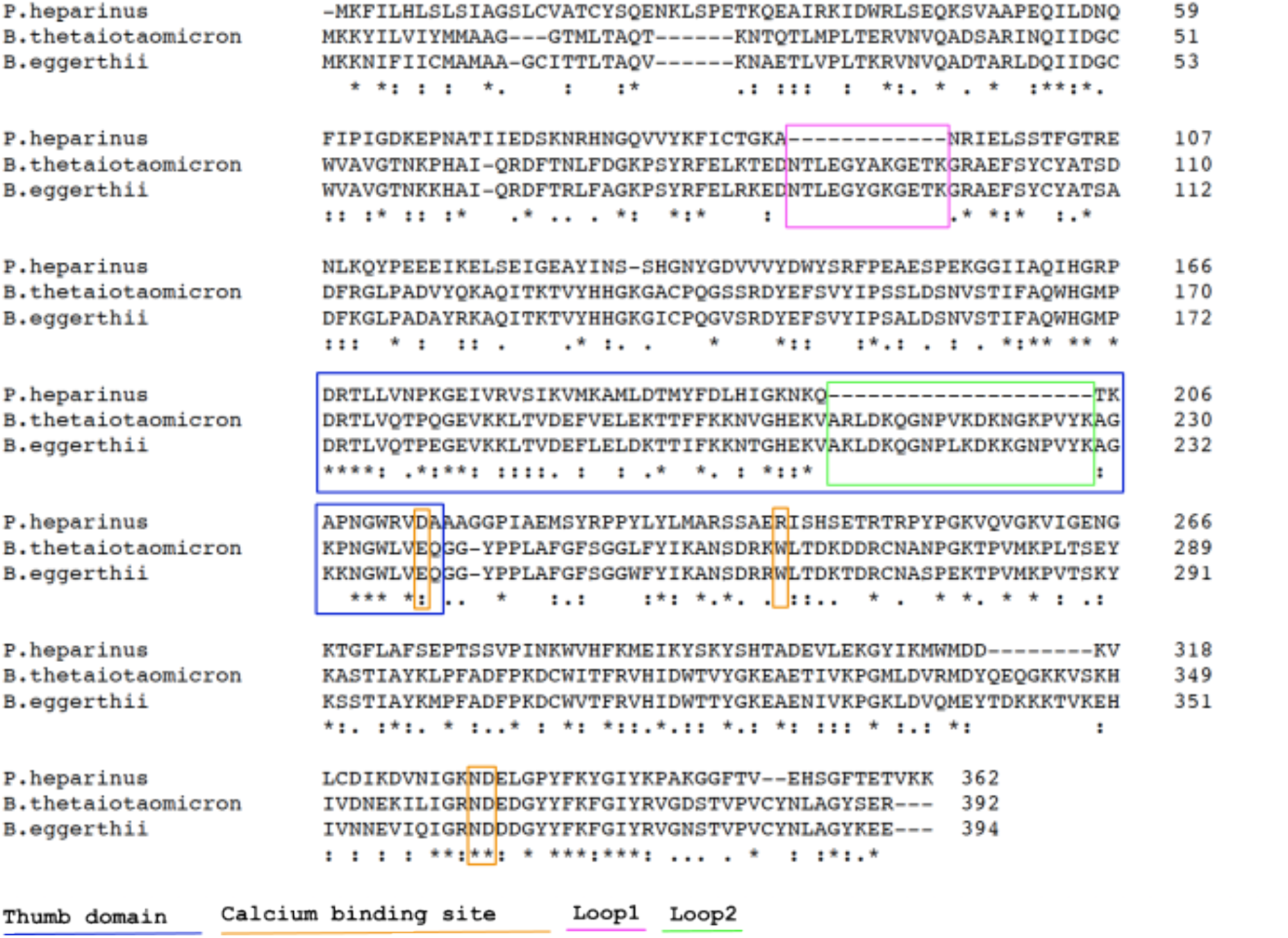
Multiple sequence alignment of *Be*HepI, *Bt*Hep1 and *Ph*HepI.

**Supplementary Figure 7.**
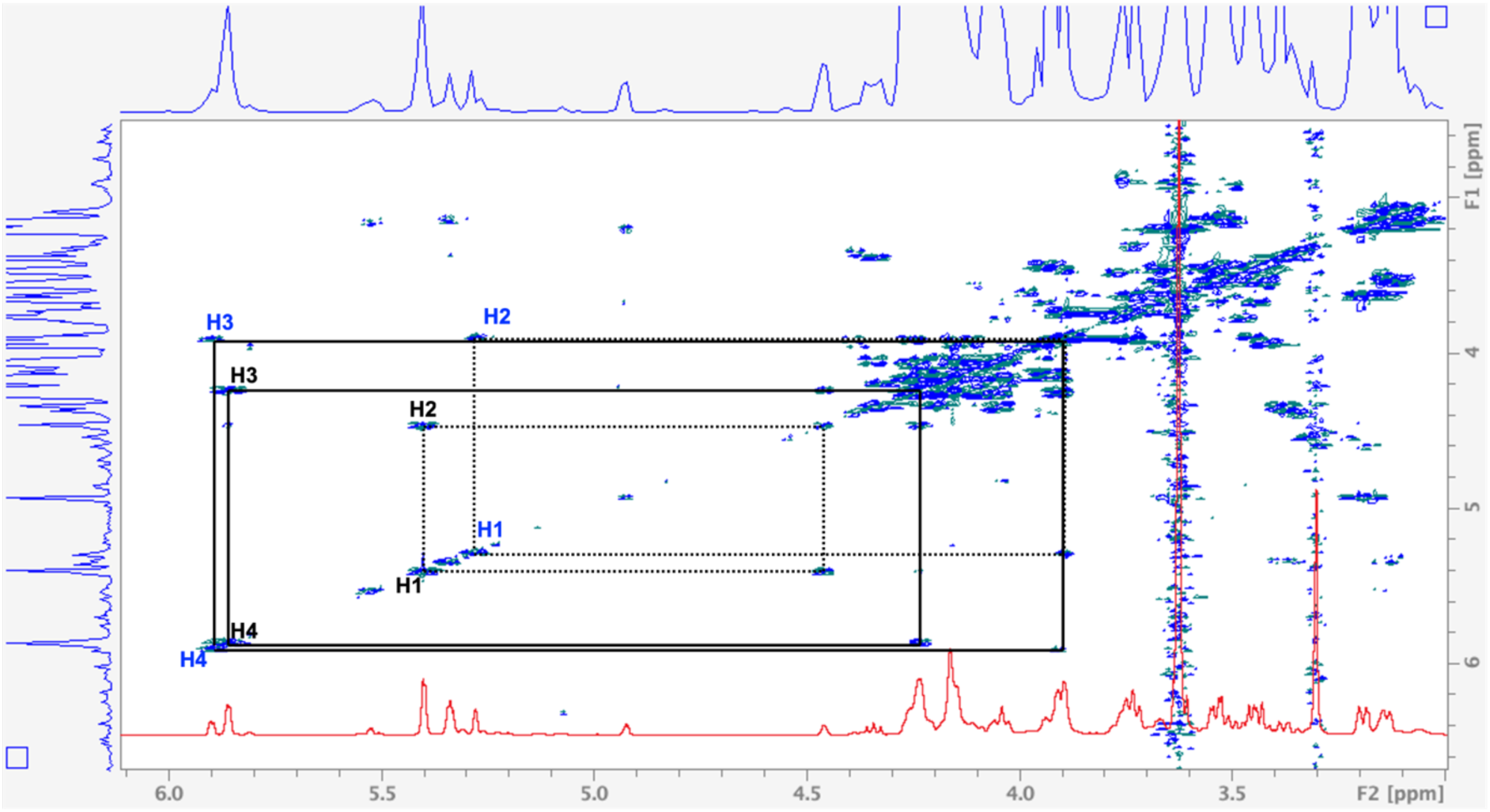
COSY NMR spectrum of fondaparinux sodium following digestion with BeHepI (ΔU2S and ΔU)

**Supplementary Figure 8.**
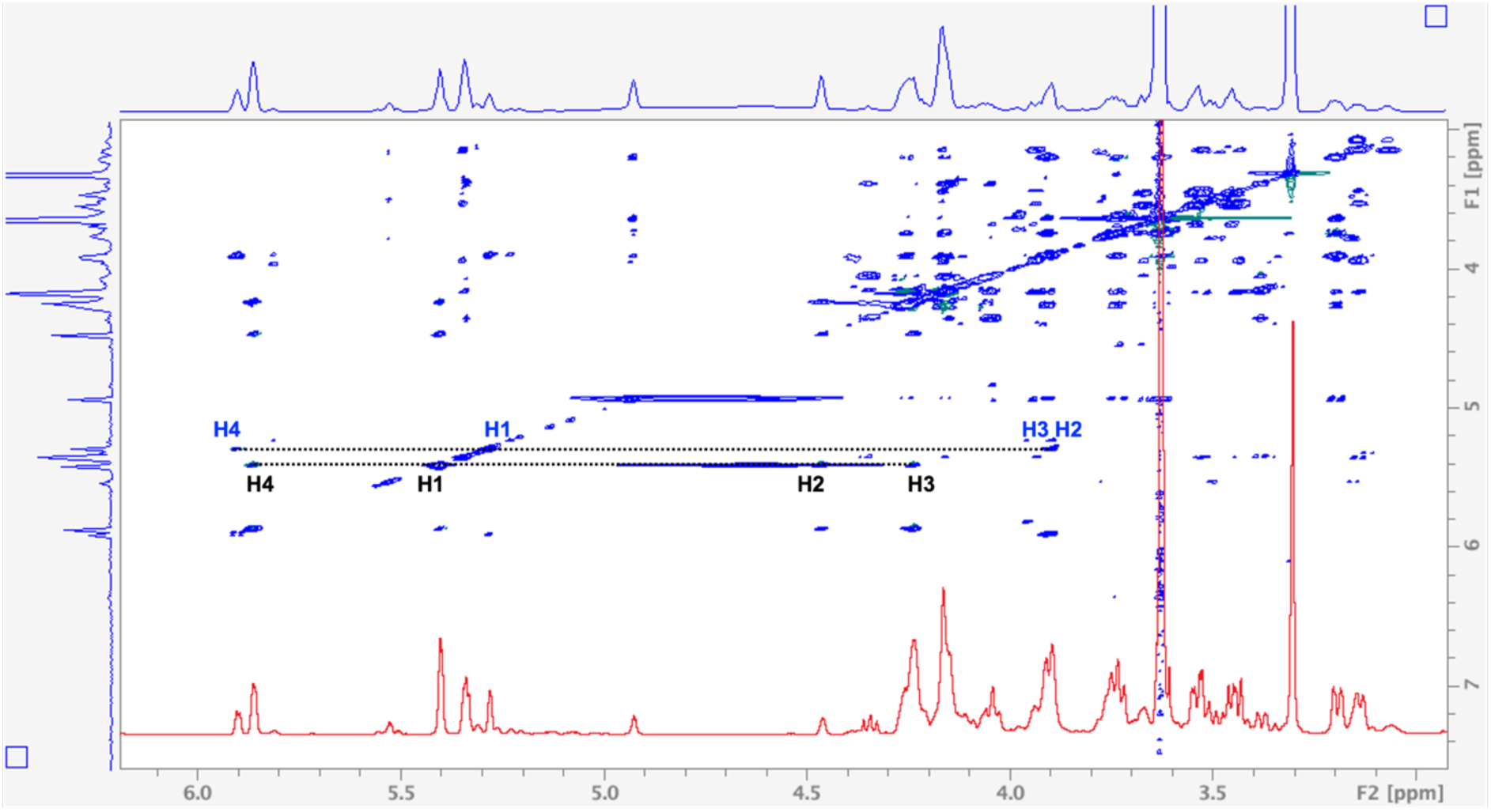
TOCSY NMR spectrum of fondaparinux sodium following digestion with *Be*HepI (ΔU2S and ΔU)

**Supplementary Figure 9.**
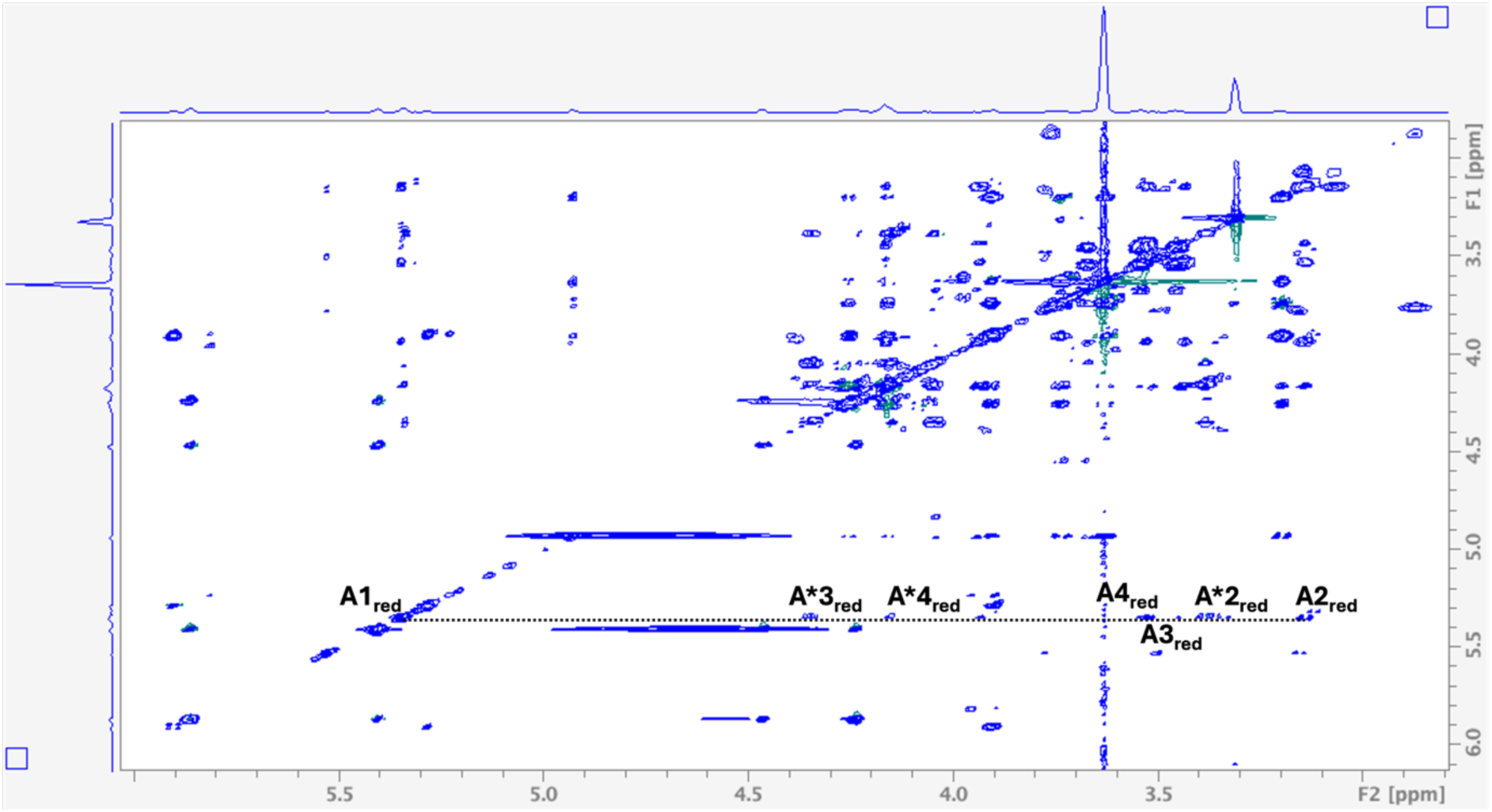
TOCSY NMR spectrum of fondaparinux sodium following digestion with *Be*HepI. Reducing end glucosamines are labelled within the figure.

## Notes

### Competing Interest Statement

The authors have declared no competing interest.

### Summary of Updates

Amended authors list (original entry was incomplete).

## References

1. Al-Hakim A. General Considerations for Diversifying Heparin Drug Products by Improving the Current Heparin Manufacturing Process and Reintroducing Bovine Sourced Heparin to the US Market. Clinical and Applied Thrombosis/Hemostasis. 2021;27. doi:10.1177/10760296211052293

2. Atha, H.D.; Lormeau, C.J., Petitou, M.; Rosenberg, D.R.; Choay, J.; Contribution of monosaccharide residues in heparin binding to antithrombin III. Biochemistry (1985), 24(23): 6723–6729. 10.1021/bi00344a063

3. Bielik, A., McLeod, E., Landry, D., Mendoza, V., Zaia, J. and Guthrie, E., 2011, November. Characterisation of Three Novel Heparinases Cloned from Bacteroides Eggerthii. In Annual Conference of the Society for Glycobiology.

4. Björk, I.; Lindahl, U.; Mechanism of the anticoagulant action of heparin. Molecular and Cellular Biochemistry.48:161–182

5. Boyce, A, and Walsh, G., 2022. Production, characteristics, and applications of microbial heparinases. Biochimie, 198, pp.109–140.

6. Choay, J.; Lormeau, J.C.; Petitou, M.; Sinaÿ, P.; Casu, B.; Oreste, P.; Torri, G.; Gatti, G.; Anti-Xa active heparin oligosaccharides. Thrombosis Research. 1980, 18 (3-4):573–578.10.1016/0049-3848(80)90356-4.

7. Chopra, P.; Joshi, A.; Wu, J.; Boons, G.J.; The 3-*O*-sulfation of heparan sulfate modulates protein binding and lyase degradation. PNAS. 2021, 118(3).10.1073/pnas.2012935118.

8. Christianson, C.H.; Belting, M.; Heparan sulfate proteoglycan as a cell-surface endocytosis receptor. Matric Biology. 35:51–55. 10.1016/j.matbio.2013.10.004.

9. Collins, E.L.; Troeberg, L.; Heparan sulfate as a regulator of inflammation and immunity. J Leukoc biol. 2019, 105:81–92. DOI:10.1002/JLB.3RU0618-246R.

10. Córdula, R.C; Lima, A.M.; Shinjo, K.S.; Gesteira, F.T.; Pol-Fachin, L.; Coulson-Thomas, J.V.; Verli, H.; Yates, A.E.; Rudd, R.T.; Pinhal, S.; Toma, L.; Dietrich, P.C.; Nader, B.H.; Tersariol, S.L.I.; On the catalytic mechanism of polysaccharide lyases: evidence of His and Tyr involvement in heparin lysis by heparinase I and the role of Ca^2+^. Molecular BioSystems. 2014, 10:54–64. DOI 10.1039/C3MB70370C.

11. David, J. G.; Wilson, S. K.; Henrissat, B. Nomenclature for sugar-binding subsites in glycosyl hydrolases. Biochem J (1997), 321(2):557–559. doi: 10.1042/bj3210557.

12. Desai, R.U.; Hui-ming, W.; Linhardt, J.R.; Specificity Studies on the Heparin Lyases from *Flavobacterium heparinum*. Biochemistry. 1993, 32:8140–8145.

13. Devlin, A.; Mycroft-West, C.; Procter, P.; Cooper, L.; Guimond, S.; Lima, M.; Yates, E.; Skidmore, M.; Tools for the Quality Control of Pharmaceutical Heparin. Medicina. 2019, 25(55):636. doi: 10.3390/medicina55100636.

14. Ding, Yili, Vara Prasad, Chamakura V. N. S., Wang, Bingyun, Advances in Chemical Synthesis of Fondaparinux, Journal of Chemistry, 2019, 9545297, 17 pages, 2019. 10.1155/2019/9545297

15. Han, H.Y.; Garron, L.M.; Kim, Y.H,; Kim, S.W.; Zhang, Z.; Ryu, S.K.; Shaya, D.; Xiao, Z.; Cheong, C.; Kim, S.Y.; Linhardt, J.R.; Jeon, H.Y.; Cygler, M. Structural snapshots of Heparin Depolymerization by Heparin Lyase I. J. Biol Chem. 2009, 284 (49):34019–34027. doi: 10.1074/jbc.M109.025338.

16. Guerrini, M.; Guglieri, S.; Naggi, A.; Sasisekharan, R.; Torri, G.; Low Molecular weight Heparins: Structural Differentiation by Bidimensional Nuclear Magnetic Resonance Spectroscopy. Semin Thromb Hemost. 2007, 33(5)L 478–48. DOI: 10.1055/s-2007-982078.

17. Guimond, E.S.; Turnbull E.J. Fibroblast growth factor receptor signalling is dictated by specific heparan sulphate saccharides. Current Biology. 1999, 9(22):1343–1346. 10.1016/S0960-9822(00)80060-3.

18. Hemker, H.C.; A century of heparin:past, present, and future. J. Throm. Haemost. 2016, 14(12):2329–2338. 10.1111/jth.13555.

19. Hogwood, J.; Mulloy, B.; Lever, R.; Grey, E.; Page, P.P.; Daws, L.; Pharmacology of Heparin and Related Drugs: An Update. Pharmacological Reviews. 2023, 75 (2):328–379. DOI: 10.1124/pharmrev.122.000684

20. Huang, Y., Mao, Y., Zong, C., Lin, C., Boons, G.J. and Zaia, J., 2015. Discovery of a heparan sulfate 3-O-sulfation specific peeling reaction. Analytical chemistry, 87(1), pp.592–600.

21. Lima, A.M.; Hughes, J.A.; Veraldi, N.; Rudd, R.T.; Hussain, R.; Brito, S.A.; Chavante, F.S.; Tersariol, I.I.; Siligardi, G.; Nader, B.H.; Yates, A.E.; Antithrombin stabilization by sulfated carbohydrates correlates with anticoagulant activity. *Med*. Chem. Commun. 2013, 4:870–873: DOI 10.1039/C3MD00048F

22. Lindahl, U.; Bäckström, G.; Höök, M.; Thunberg, L.; Fransson, L.; Linker, A.; Structure of the antithrombin-binding site in heparin. Proc. Natl. Acad. Sci. USA. 1979, 76 (7):3198–3202. 10.1073/pnas.76.7.3198

23. Liu, Y-C.; Su, B-W.; Guo, L-B.; Zhang Y-W.; Cloning, expression, and characterisation of a novel heparinase I from *Bacteroides eggerthii*. Preparative Biochemistry & Biotechnology. 2020, 50:477–485.10.1080/10826068.2019.1709977.

24. Meneghetti, C.Z.M.; Hughes, J.A.; Rudd, R, T.; Nader, B.H.; Powell, K.A.; Yates, A.E.; Lima, A.M. Heparan sulfate and heparin interactions with proteins. J. R. Soc. Interface. 2015, 12:10150589.10.1098/rsif.2015.0589.

25. Olson, T.S.; Björk, I.; Sheffer, R.; Craig, A.P.; Shore, D.J.; Choay, J.; Role of the Antithrombin-binding Pentasaccharide in Heparin Acceleration of Antithrombin-Proteinase Reactions. J. Biol Chem.1992, 267 (18):12528–12538.

26. Tovar, M.A.; Santos, C.R.G.; Capillé, V.N.; Piquet, A.A.; Glauser, F.B.; Pereira, S.M.; Vilanova, E.; Mourão, S.A.P.; Structural and haemostatic features of pharmaceutical heparins from different animal sources: challenges to define thresholds separating distinct drugs. Scientific reports: (2016), 6, 35619. 10.1038/srep35619

27. Ravikumar, M.; Smith, A. A.R; Nurcombe, V.; Cool, M.S.; Heparan Sulfate Proteoglycans: Key Mediators of Stem Cell Function. Front. Cell. Biol. 2020, 8:DOI:https://doi,org/10.3389/fcell.2020.581213.

28. Rosenberg, D.R.; Lam, L.; Correlation between structure and function of heparin. PNAS. 1979, 76(3):1218–1222. 10.1073/pnas.76.3.1218.

29. Shriver, Z.; Sundaram, M.; Venkataraman, G.; Fareed, J.; Linhardt, R.; Biemann, K.; Sasisekharan, R.; Cleavage of the antithrombin III binding site in heparin by heparinases and its implication in the generation of low molecular weight heparin. PNAS. 2000, 97(19):10365–10370. 10.1073/pnas.97.19.10365.

30. Shukla, D.; Liu. J.; Blaiklock, P.; Shworak, W.N.; Bai, X.; Esko, D.J.; Chohen, H.G.; Eisenberg, J.R.; Rosenberg, D.R.; Spear, G.P.; A Novel Role for 3-O-sulphated Heparan sulfate in Herpes Simplex Virus 1 Entry. Cell. 1999, 99(1):13–22. 10.1016/S0092-8674(00)80058-6.

31. Yamada, S.; Yoshida, K.; Sugiura, M.; Sugahara, K.; Khoo, H.K.; Morris, R.H.; Dell, A.; Structural studies on the bacterial lyase resistant tetrasaccharide derived from the antithrombin III-binding site of porcine intestinal heparin. Jornal of Biological Chemistry. 1993, 268(7):4780–4787. 10.1016/S0021-9258(18)53465-7.

32. Zhao, W.; Garron, L-M.; Yang, B.; Xiao, Z.; Esko, D.J.; Cygler, M.; Linhardt, J.R.; Asparagine 405 of heparin lyase II prevents the cleavage of glycosidic linkages proximate to a 3-O-sulfoglucosamine residue. FEBS LettersI.2011, 585(25):2461–2466. 10.1016/j.febslet.2011.06.023.

